# Towards reconstructing intelligible speech from the human auditory cortex

**DOI:** 10.1101/350124

**Authors:** Hassan Akbari, Bahar Khalighinejad, Jose L. Herrero, Ashesh D. Mehta, Nima Mesgarani

## Abstract

Auditory stimulus reconstruction is a technique that finds the best approximation of the acoustic stimulus from the population of evoked neural activity. Reconstructing speech from the human auditory cortex creates the possibility of a speech neuroprosthetic to establish a direct communication with the brain and has been shown to be possible in both overt and covert conditions. However, the low quality of the reconstructed speech has severely limited the utility of this method for brain-computer interface (BCI) applications. To advance the state-of-the-art in speech neuroprosthesis, we combined the recent advances in deep learning with the latest innovations in speech synthesis technologies to reconstruct closed-set intelligible speech from the human auditory cortex. We investigated the dependence of reconstruction accuracy on linear and nonlinear (deep neural network) regression methods and the acoustic representation that is used as the target of reconstruction, including auditory spectrogram and speech synthesis parameters. In addition, we compared the reconstruction accuracy from low and high neural frequency ranges. Our results show that a deep neural network model that directly estimates the parameters of a speech synthesizer from all neural frequencies achieves the highest subjective and objective scores on a digit recognition task, improving the intelligibility by 65% over the baseline method which used linear regression to reconstruct the auditory spectrogram. These results demonstrate the efficacy of deep learning and speech synthesis algorithms for designing the next generation of speech BCI systems, which not only can restore communications for paralyzed patients but also have the potential to transform human-computer interaction technologies.

## Introduction

Auditory stimulus reconstruction is an inverse mapping technique that finds the best approximation of the acoustic stimulus from the population of evoked neural activity. Stimulus reconstruction was originally proposed as a method to study the representational properties of the neural population ^1–5^ because this method enables the intuitive interpretation of the neural responses in the stimulus domain. Reconstructing speech from the neural responses recorded from the human auditory cortex^6^, however, opens up the possibility of using this technique as a speech brain-computer interface (BCI) to restore speech in severely paralyzed patients (for a review, see these references^7–9^). The ultimate goal of a speech neuroprosthesis is to create a direct communication pathway to the brain with the potential to benefit patients who have lost their ability to speak, which can result from a variety of clinical disorders leading to conditions such as locked-in syndrome^10,11^. The practicality of using speech decoding methods in a neuroprosthetic device to restore speech communication was further supported by studies showing successful decoding of speech during both overt and covert (imagined) conditions ^12–16^. These studies showed successful decoding of imagined articulations^13,14^, imagined word repetition^15^, and silent reading of speech^16^ from auditory cortical areas, including the superior temporal gyrus (STG). While previous studies have established the feasibility of reconstructing speech from neural data, the quality of the reconstructed audio so far has been too low to merit subjective evaluation. For this reason, the reconstructed sounds in previous studies have been evaluated only using objective measures such as correlation or recognition accuracy^3,6,8,13,17–25^. The low quality of the reconstructed sound is currently a major limiting factor in actualizing speech BCI systems^7^.

The acoustic representation of the stimulus that is used as the decoding target can significantly impact the quality and accuracy of reconstructed sounds. Previous studies have used magnitude spectrogram (time-frequency representation)^3,20^, speech envelope^21,22^, spectrotemporal modulation frequencies^6,13,23^, and discrete units such as phonemes and phonetic categories^8,17,24,25^ and words^18,19^. Using discrete units can be advantageous by allowing for discriminative training. However, decoding discrete representations of speech such as phonemes eliminates the paralinguistic information such as speaker features, emotion, and intonation. In comparison, reconstructing continuous speech provides the possibility of real-time, continuous feedback that can be delivered to the user to promote coadaptation of the subject and the BCI algorithm^26,27^ for enhanced accuracy. A natural choice is to directly estimate the parameters of a speech synthesizer from neural data, but this has not been attempted previously because the process requires a highly accurate estimation of several vocoder parameters, which is hard to achieve with traditional machine-learning techniques.

To advance the state-of-the-art in speech neuroprosthesis, we aimed to increase the intelligibility of the reconstructed speech by combining recent advances in deep learning^28^ with the latest innovations in speech synthesis technologies. Deep learning models have recently become the dominant technique for acoustic and audio signal processing^29–32^. These models can improve reconstruction accuracy by imposing more complete constraints on the reconstructed audio by better modeling the statistical properties of the speech signal^3^. At the same time, nonlinear regression can invert the nonlinearly encoded speech features in neural data^33,34^ more accurately.

We examined the effect of three factors on the reconstruction accuracy: 1) the regression technique (linear regression versus nonlinear deep neural network), 2) the representation of the speech intended for reconstruction (auditory spectrogram versus speech vocoder parameters), and 3) the neural frequency range used for regression (low frequency versus high-gamma envelope) (Fig. 1A). Our results showed that a deep neural network model that uses all neural frequencies to directly estimate the parameters of a speech vocoder achieves the highest subjective and objective scores, both for intelligibility and the quality of reconstruction in a digit recognition task. These results represent an important step toward successful implementation of the next generation of speech BCI systems.

## Results

### Neural recordings

We used invasive electrocorticography (ECoG) to measure neural activity from five neurosurgical patients undergoing treatment for epilepsy as they listened to continuous speech sounds. Two of the five subjects had high-density subdural grid electrodes implanted in the left hemisphere with coverage primarily over the superior temporal gyrus (STG), and four of the five subjects had depth electrodes with coverage of Heschl's gyrus (HG). All subjects had self-reported normal hearing. Subjects were presented with short continuous stories spoken by four speakers (two females, total duration: 30 minutes). To ensure that the subjects were engaged in the task, the stories were randomly paused, and the subjects were asked to repeat the last sentence.

The test data consisted of continuous speech sentences and isolated digit sounds. We used eight sentences (40 seconds total) to evaluate the objective quality of the reconstruction models. The sentences were repeated six times in random order, and the neural data was averaged over the six repetitions to reduce the effect of neural noise on comparison of reconstruction models (see Supp. Fig. 1 for the effect of averaging). The digit sounds were used for subjective intelligibility and quality assessment of reconstruction methods and were taken from a publicly available corpus, TI-46^35^. We chose 40 digit sounds (zero to nine), spoken by four speakers (two females) that were not included in the training of the models. Reconstructed digits were used as the test set to evaluate subjective intelligibility and quality of the models. Two ranges of neural frequencies were used in the study. Low-frequency (0-50 Hz) components of the neural data were extracted by filtering the neural signals using a lowpass filter. The high-gamma envelope^36^ was extracted by filtering the neural signals (70 to 150 Hz) and calculating the Hilbert envelope^37^.

### Regression models

The input to the regression models was a sliding window over the neural data with a duration of 300 ms (Fig. 1B), and the hop size of 10 ms. The duration of the sliding window was chosen to maximize reconstruction accuracy (supp. Fig. 2). We compared the performance of linear and nonlinear regression models to reconstruct the stimulus from the neural signals. The linear regression finds a linear mapping between the response of a population of neurons to the stimulus representation^3,6^. This method effectively assigns a spatiotemporal filter to each electrode estimated by minimizing the mean-squared-error (MSE) between the original and reconstructed stimulus.

The nonlinear regression model was implemented using a deep neural network (DNN). We designed a deep neural network architecture with two stages: 1) feature extraction and 2) feature summation networks^38–40^ (Fig. 1B). In this framework, a high-dimensional representation of the input (neural responses) is first calculated, which results in mid-level features (output of the feature extraction network). These mid-level features are then input to the feature summation network to regress the output of the model (acoustic representation). The feature summation network in all cases was a two-layer fully connected network (FCN) with regularization, dropout^41^, batch normalization^42^, and nonlinearity between each layer. For feature extraction, we compared the efficacy of five different network architectures for auditory spectrogram and vocoder reconstruction (Methods, Supp. Table 1 for details of each network). Specifically, we found that the fully connected network (FCN), in which no constraint was imposed on the connectivity of the nodes in each layer of the network to the previous layer, achieved the best performance for reconstructing the auditory spectrogram. However, the combination of the FCN and a locally connected network (LCN), which constrains the connectivity of each node to only a subset of nodes in the previous layer, achieved the highest performance for the vocoder representation (Supp. Tables 4, 5). In the combined FCN+LCN, the outputs of the two parallel networks are concatenated and used as the mid-level features (Fig.1B).

**Figure 1.**
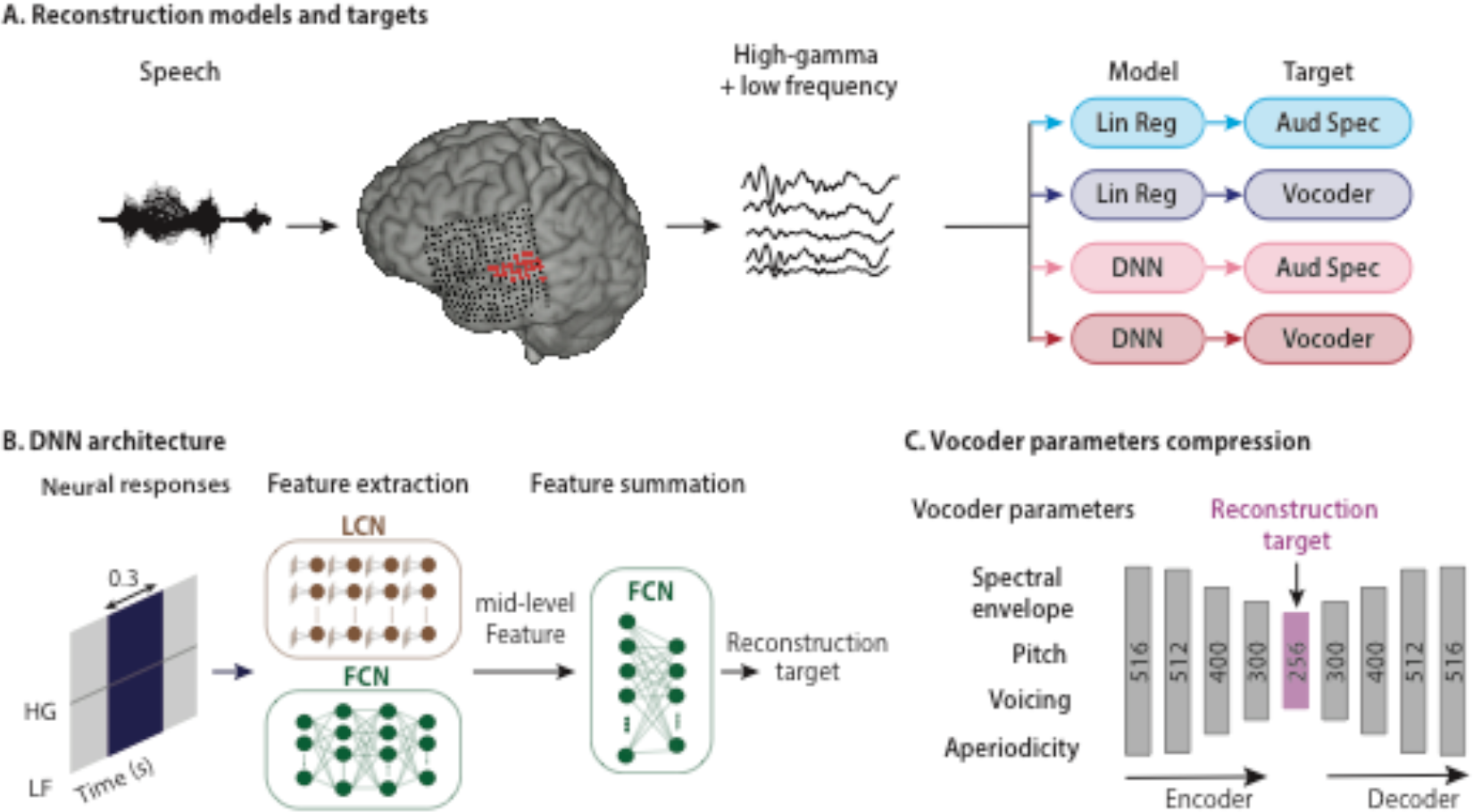
Schematic of the speech reconstruction method. (A) Subjects listened to natural speech sentences. The population of evoked neural activity in the auditory cortex of the listener was then used to reconstruct the speech stimulus. The responsive electrodes in an example subject are shown in red. High and low frequency bands were extracted from the neural data. Two types of regression models and two types of speech representations were used, resulting in four combinations: linear regression to auditory spectrogram (light blue), linear regression to vocoder (dark blue), DNN to auditory spectrogram, and DNN to vocoder (dark red). (B) The input to all models was a 300 ms sliding window containing both low frequency (LF) and the high-gamma envelope (HG). The DNN architecture consists of two modules: feature extraction and feature summation networks. Feature extraction for auditory spectrogram reconstruction was a fully connected neural network (FCN). For vocoder reconstruction, the feature extraction network consisted of an FCN concatenated with a locally connected network (LCN). The feature summation network is a two-layer fully connected neural network (FCN). (C) Vocoder parameters consist of spectral envelope, fundamental frequency (*f*0), voicing, and aperiodicity (total of 516 parameters). An autoencoder with a bottleneck layer was used to reduce the 516 vocoder parameters to 256. The bottleneck features were then used as the target of reconstruction algorithms. The vocoder parameters were calculated from the reconstructed bottleneck features using the decoder part of the autoencoder network.

### Acoustic representations

We used two types of acoustic representation of the audio as the target for reconstruction: auditory spectrogram and speech vocoder. The auditory spectrogram was calculated using a model of the peripheral auditory system^43,44^, which estimates a time-frequency representation of the acoustic signal on a tonotopic frequency axis. The reconstruction of the waveform from the auditory spectrogram is achieved using an iterative convex optimization procedure^43^ because the phase of the signal is lost during this procedure.

For speech vocoder, we used a vocoder-based, high-quality speech synthesis algorithm (WORLD^45^), which synthesizes speech from four main parameters: 1) spectral envelope, 2) *f0* or fundamental frequency, 3) band aperiodicity, and 4) a voiced-unvoiced (VUV) excitation label (Fig. 1C). These parameters are then used to re-synthesize the speech waveform. This model can reconstruct high-quality speech and has been shown to outperform other methods including STRAIGHT^46^. The large numbers of the parameters in the vocoder (516 total) and the susceptibility of the synthesis quality on inaccurate estimation of parameters however pose a challenge. To remedy this, we first projected the sparse vocoder parameters onto a dense subspace in which the number of parameters can be reduced, which allows better training with a limited amount of data. We used a dimensionality reduction technique that relies on an autoencoder (AEC)^47^(Fig. 1C), which compresses the vocoder parameters into a smaller space (encoder, 256 dimensions, Supp. Table 3) and subsequently recovers (decoder) the original vocoder parameters from the compressed features (Fig. 1C). The compressed features (also called bottleneck features) are used as the target for the reconstruction network. By adding noise to the bottleneck features before feeding them to the decoder during training, we can make the decoder more robust to unwanted variations in amplitude, which is necessary due to the noise inherently present in the neural signals. The autoencoder was trained on 80 hours of speech using a separate speech corpus (Wall Street Journal l^48^). During the test phase, we first reconstructed the bottleneck features from the neural data, and subsequently estimated the vocoder parameters using the decoder part of the autoencoder (Fig. 1C). The reconstruction accuracy of individual vocoder parameters with a neural network shows varied improvement over the linear model, where pitch estimation is improved the most (%157.2), followed by aperiodicity (%18.5), spectral envelope (%6.2), and voiced-unvoiced parameter (%0.15, Supp. Fig. 3).

Figure 2B shows the example reconstructed auditory spectrograms from each of the four combinations of the regression models (linear regression and DNN) and acoustic representation (auditory spectrogram and vocoder). Comparison of the auditory spectrograms in Figure 2A shows that 1) the overall frequency profile of the speech utterance is better preserved by the DNN compared to the linear regression model, and 2) the harmonic structure of speech is recovered only in the DNN-Vocoder model.

These observations are shown more explicitly in Figure 2B, where the magnitude power of frequency bands is shown during an unvoiced (t = 1.4 sec) and a voiced speech sound (t = 1.15 sec, shown with dashed lines). The frequency profile of original and reconstructed auditory spectrograms during the unvoiced sound shows a more accurate reconstruction of low and high frequencies for the DNN models (Fig. 2B left, comparison of blue and red plots). The comparison of frequency profiles during the voiced sound (Fig. 2B, right) reveals the recovery of speech harmonics only in the DNN-Vocoder model (comparison of top and bottom plots).

**Figure 2.**
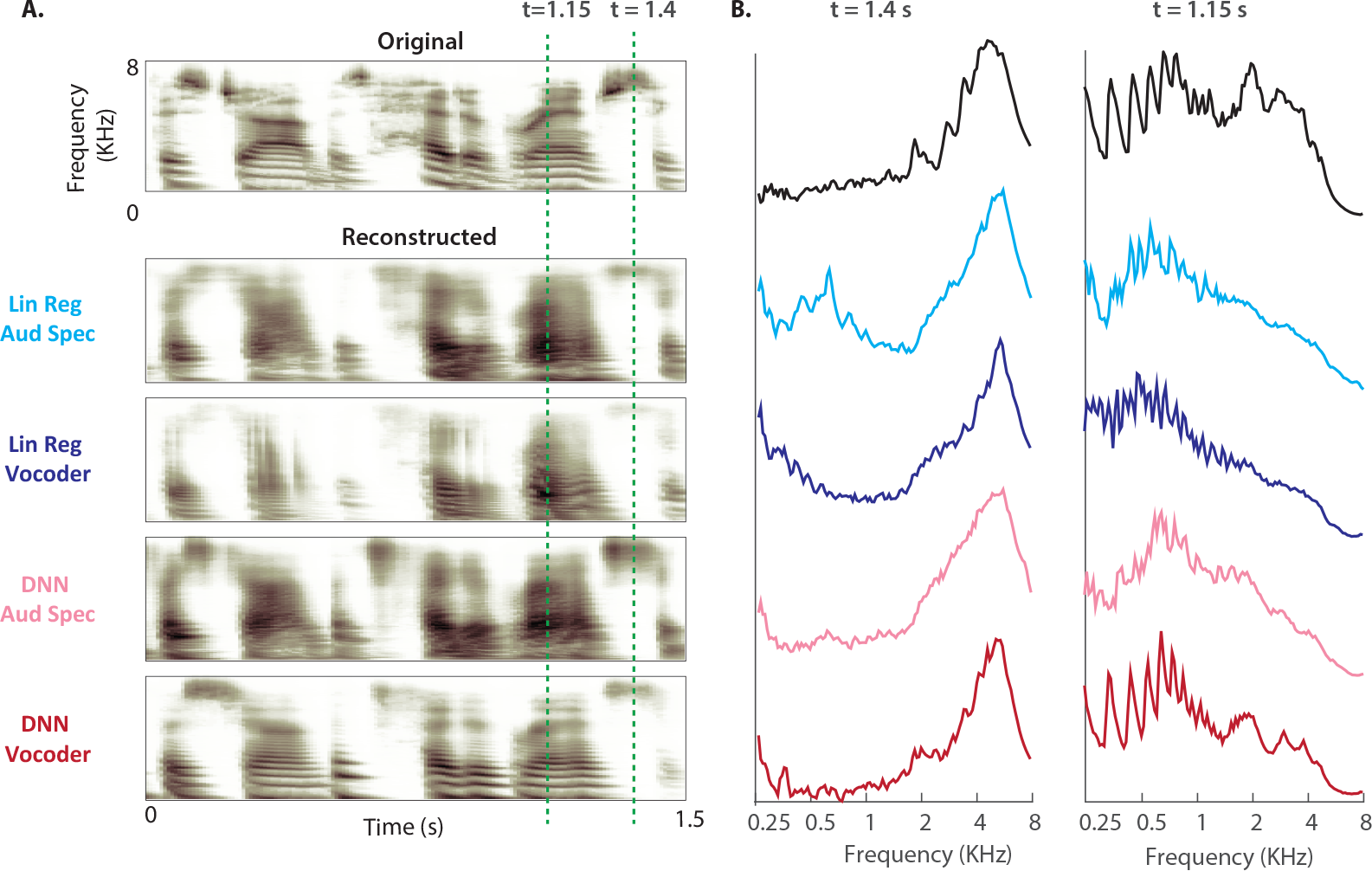
Deep neural network architecture. (A) An original auditory spectrogram of a speech sample is shown on top. The reconstructed auditory spectrograms of the four models are shown below. (B) Magnitude power of frequency bands during an unvoiced (t = 1.4 sec) and a voiced speech sound (t = 1.15 sec, shown with dashed lines in A) for original (top) and the four reconstruction models.

### Subjective evaluation of the reconstruction accuracy

We used the reconstructed digit sounds to assess the subjective intelligibility and quality of the reconstructed audio. Forty unique tokens were reconstructed from each model, consisting of ten digits (zero to nine) that were spoken by two male and two female speakers. The speakers that uttered the digits were different from the speakers that were used in the training, and no digit sound was included in the training of the networks. We asked 11 subjects with normal hearing to listen to the reconstructed digits from all four models (160 tokens total) in a random order. Each digit was heard only once. The subjects then reported the digits (zero to nine, or uncertain), rated the reconstruction quality using the mean opinion score (MOS^49^, on a scale of 1 to 5), and reported the gender of the speaker (Fig. 3A).

Figure 3B shows the average reported intelligibility of the digits from the four reconstruction models. The DNN-vocoder combination achieved the best performance (75% accuracy), which is 67% higher than the baseline system (Linear regression with auditory spectrogram). Fig. 3B also shows that the reconstructions using DNN models are significantly better than the linear regression models (68.5% vs. 47.5%, paired t-test, p<0.001). Figure 3C shows that the subjects also rated the quality of the reconstruction significantly higher for the DNN-vocoder system than for the other three models (3.4 vs. 2.5, 2.3, and 2.1, unpaired t-test, p<0.001), meaning that the DNN-vocoder system sounds closest to natural speech. The subjects also accurately reported the gender of the speaker significantly higher than chance for the DNN-vocoder system (80%, t-test, p<0.001) while the performance for all other methods were at chance (Fig. 3D). The higher intelligibility and quality scores for the DNN-Voc system was consistently observed in all the ten listeners (Supp. Fig. 4). This result indicates the importance of accurate reconstruction of harmonics frequencies for identifying speaker dependent information, which are best captured by the DNN-Voc model.

Finally, Figure 3E shows the confusion patterns in recognizing the digits for the four models, confirming again the advantage of the DNN based models, and the DNN vocoder in particular. As shown in Figure 3E, the discriminant acoustic features of the digit sounds are better preserved in the DNN-Voc model, enabling the listeners to correctly differentiate them from the other digits. Linear regression models, however, failed to preserve these cues, as seen by the high confusion among digit sounds. The confusion patterns also show that some errors were associated with the shared phonetic features, for example the confusion between digits one and nine (sharing ‘ey’ phoneme), or four and fine (sharing the initial fricative /f/ phoneme. This result suggests a possible strategy for enabling accurate discrimination in BCI applications by selecting target sounds with a sufficient acoustic distance between them. The audio samples from different models can be found online^50^ and in the supplementary materials.

**Figure 3.**
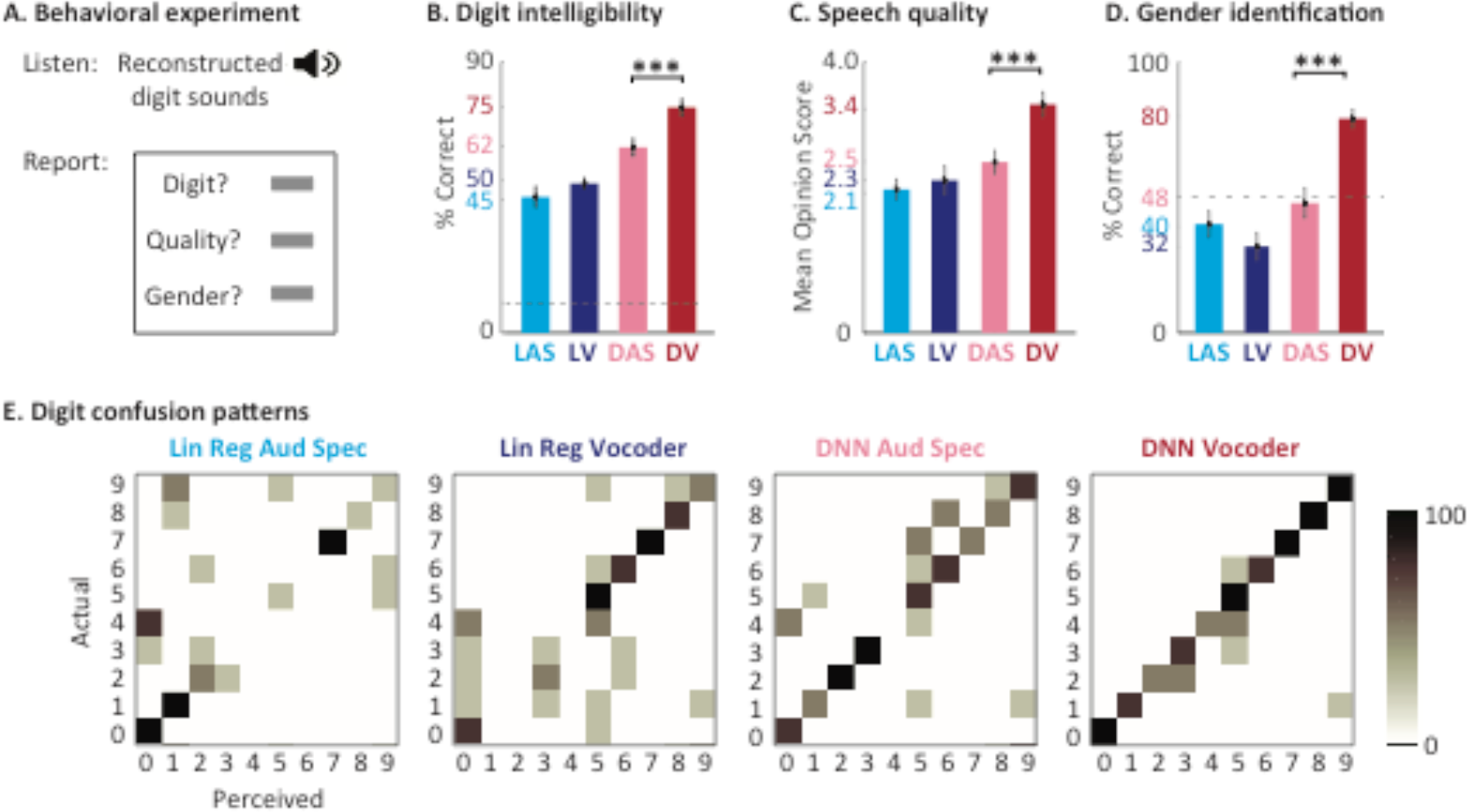
Subjective evaluation of the reconstruction accuracy. (A) The behavioral experiment design used to test the intelligibility and the quality of the reconstructed digits. Eleven subjects listened to digit sounds (zero to nine) spoken by two male and two female speakers. The subjects were asked to report the digit, the quality on the mean-opinion-scale, and the gender of the speaker. (B) The intelligibility score for each model defined as the percentage of correct digits reported by the subject. (C) The quality score on the MOS scale. (D) The speaker gender identification rate for each model. (E) The digit confusion patterns for each of the four models. The DNN vocoder shows the least amount of confusion among the digits.

### Objective evaluation of reconstructed audio

We compared the objective reconstruction accuracy of reconstructed audio per subject using the extended short time objective intelligibility (ESTOI)^51^ measure. ESTOI is commonly used for the intelligibility assessment of speech synthesis technologies and is calculated by measuring the distortion in spectrotemporal modulation patterns of the noisy speech signal. Therefore, ESTOI score is sensitive to both inaccurate reconstruction of the spectral profile and the inconsistencies in the reconstructed temporal patterns. The ESTOI measures were calculated from continuous speech sentences in the test set. The average ESTOI of the reconstructed speech for all five subjects (Fig. 4A) confirms the results seen from the subjective tests, which is the superiority of DNN based models over the linear model, and that of vocoder reconstruction over the auditory spectrogram (p<0.001, t-test). This pattern was consistent for each of the five subjects in this study, as shown in Fig. 4B alongside the electrode locations for each subject. While the overall reconstruction accuracy varies significantly across subjects, which is likely due to the difference in the coverage of the auditory cortical areas, the relative performance of the four models was the same in all subjects. In addition, averaging the neural responses over multiple repetitions of the same speech utterance improved the reconstruction accuracy (Supp. Fig. 1) because averaging reduces the effect of neural noise.

**Figure 4.**
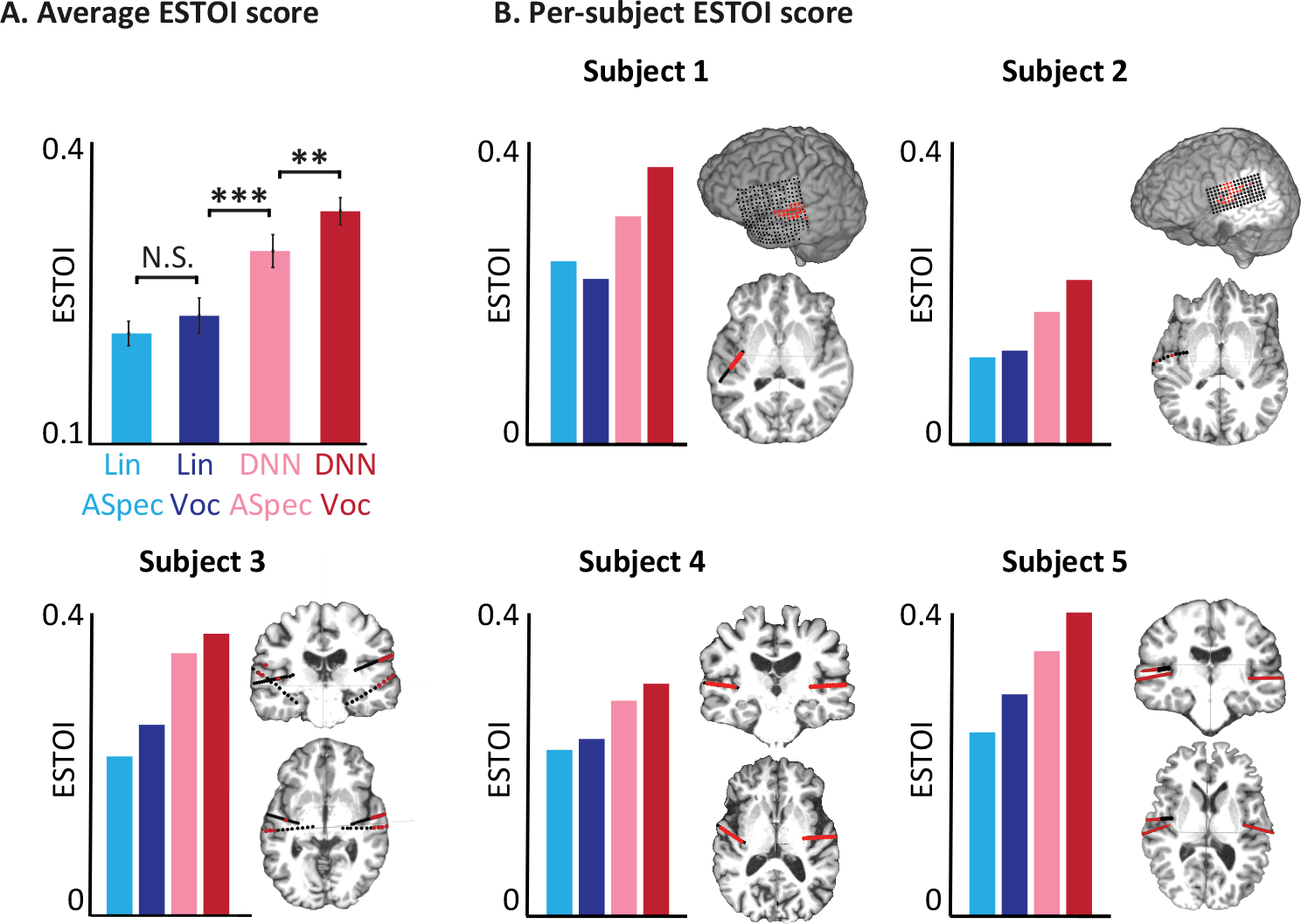
Objective intelligibly scores for different models. (A) The average ESTOI score based on all subjects for the four models. (B) Coverage and the location of the electrodes and ESTOI score for each of the five subjects. In all subjects, the ESTOI score of the DNN vocoder was higher than in the other models.

### Reconstruction accuracy from low and high neural frequencies

There is increasing evidence that the low and high-frequency bands encode different and complementary information about the stimulus^52^. Considering that the sampling frequency of the reconstruction target is 100 Hz, we used 0-50 Hz as a low-frequency signal, and the envelope of high gamma (70-150 Hz) as high-frequency band information. To determine what frequency bands are best to include to achieve maximum reconstruction accuracy, we tested the reconstruction accuracy in three conditions, when the regression model uses only the high-gamma envelope, a low-frequency signal, or a combination of the two.

To simplify the comparison, we used only the DNN-auditory spectrogram reconstruction model. We calculated the ESTOI scores of the reconstructed speech sound using different frequency bands. We found that the combination of the two frequency bands significantly outperforms the reconstruction from only one of the frequency bands (Fig. 5A, p<0.001, t-test). This observation is consistent with the complementary encoding of the stimulus features in the low and high-frequency bands^53^, which implicates the advantage of using the entire neural signal to achieve the best performance in speech neuroprosthesis applications when it is practically possible.

### Effect of the number of electrodes and duration of training data

The variability of the reconstruction accuracy across subjects (Fig. 4B) suggests an important role of neural coverage in improving the reconstruction^3,6^ accuracy. In addition, because some of the noise signal across different electrodes is independent, reconstruction from a combination of electrodes may lead to a higher accuracy by finding a signal subspace less affected by the noise in the data^54^. To examine the effect of the number of electrodes on the reconstruction accuracy, we first combined the electrodes of all five subjects and randomly chose N electrodes (N = 1, 2, 4, 8, 16, 32, 64, 128), twenty times for training the individual networks. The average reconstruction accuracy for each N was then used for comparison. The results shown in Fig. 5B indicate that increasing the number of electrodes improves the reconstruction accuracy; however, the rate of improvement decreased significantly.

Finally, because the success of neural network models is largely attributed to training on large amounts of data^28^, we examined the effect of training duration on reconstruction accuracy. We used 128 randomly chosen electrodes and trained several neural network models each on a segment of the training data as the duration of the segments was gradually increased from 10 to 30 minutes. This process was performed twenty times for each duration by choosing a random segment of the training data, and the ESTOI score was averaged over the segments. As expected, the results show an increased reconstruction accuracy as the duration of the training was increased (Fig. 5C), which indicates the importance of collecting a larger duration of training data when it is practically feasible.

## Discussion

We compared the performance of linear and nonlinear (DNN) regression models in reconstructing the auditory spectrogram and vocoder representation of speech signals. We found that using a deep neural network model to regress vocoder parameters significantly outperformed the linear regression and auditory spectrogram representation of speech, and resulted in 75% intelligibility scores on a closed-set, digit recognition task.

Our results are consistent with those of previous reconstruction studies that showed the importance of nonlinear techniques in neural decoding^55^. The previous methods have used support vector machines^13,56^, linear discriminant analysis^57,58^, linear regression^3,14,59^, nonlinear embedding^6^, and Bayes classifiers^15^. In recent years, deep learning^60^ has shown tremendous success in many brain-computer interface technologies^61^, and our study extended this trend by showing the benefit of deep learning in speech neuroprosthesis research^55^.

We showed that the reconstruction accuracy depends on both the number of electrodes and the duration of the data that is available for training. This is consistent with the findings of studies showing the superior advantage of deep learning models over other techniques, particularly when the amount of training data is large^28^. We showed that the rate of improvement slows down as the number of electrodes increases. This could indicate the limited diversity of the neural responses in our recording which ultimately limits the added information that is gained from additional electrodes. Alternatively, increasing the number of electrodes also increases the complexity and the number of free parameters in the neural network model. Because the duration of our training data was limited, it is possible that more training data would be needed before the benefit of additional features becomes apparent. Our experiments showed that increasing the amount of training data results in better reconstruction accuracy, therefore recording methods that can increase the amount of data available for the training of deep models are highly desirable, for example, when chronic recordings are possible in long-term implantable devices such as the NeuroPace responsive neurostimulation device (RNS) ^62^.

We showed that the representation of the acoustic signal used as the target of reconstruction has an important role in the intelligibility and the quality of the reconstructed audio. We used a vocoder representation of speech, which extends the previous studies that used a magnitude spectrogram (time-frequency representation)^3,20^, speech envelope^21,22^, spectrotemporal modulation frequencies^6,13,23^, and discrete units such as phonemes and phonetic categories^8,17,24,25^ and words^18,19^. Reconstruction of the auditory spectrogram, which we also used for comparison, inherently results in suboptimal audio quality because the phase of the auditory spectrogram must be approximated. The discrete units such as phonemes enable discriminative training by learning a direct map from the neural data to the class labels, which is typically more efficient than generative regression models^63^. The continuous nature of parameters in acoustic reconstruction however could prove advantageous for BCI applications because they provide a continuous feedback to the user^64^, which is crucial for the subject and the BCI algorithm to coadapt to increase overall effectiveness^26,27^. Therefore, direct reconstruction of speech synthesis parameters is a natural choice. This choice however poses a challenge, since the vocoder quality is very sensitive to the quality of the decoding. As we have reported, reconstructing vocoder parameters resulted in both the worst (when used with linear regression) and the best (when used with DNN) results. Therefore, powerful modeling techniques such as deep learning are crucial as more inclusive representations of the speech signal are used for reconstruction and decoding applications. We proposed a solution to this problem by compressing the acoustic features into a low-dimensional space and using a decoder that is robust to the fluctuations of the input.

We found that the combination of low frequency and the envelope of high gamma results in higher reconstruction accuracy than each frequency band alone. This finding is consistent with those of studies that have shown the importance of an oscillatory phase^65^ in addition to the neural firing rate, which is reflected in the high-gamma frequency band^66^. Combining both high and low frequencies not only enables access to the complementary information in each band^52,67^ but also allows the decoder to use the information that is encoded in the interactions between the two bands, such as cross-frequency coupling^53^. Overall, we observed that better brain coverage, more training data, and combined neural frequency bands result in the best reconstruction accuracy, which can serve as an upper bound performance where practical limitations prevent the use of all possible factors, for example, where the brain coverage is small, or high-frequency neural signals are not accessible such as in noninvasive neuroimaging methods.

The application of neural speech decoding in neuroprosthesis is contingent on the similarity of the underlying neural code in overt and covert (imagined) conditions. Several previous studies have examined the generalization of decoding techniques from overt to covert speech^12–16^ and showed the involvement of the auditory cortical areas, including the superior temporal gyrus (STG) in covert speech condition. Specifically, informative electrodes for speech decoding were found in Wernicke and the STG during imagined articulation^13,14^, covert word repetition^15^, and reading silently^16^. In addition to imagined articulation, an MEG study^12^ measured the neural activity during actual and imagined hearing conditions and compared with actual and imagined articulation conditions. This study found that the neural activity during overt and covert states were more similar in hearing than in articulation condition. Furthermore, the similarity of the response topographies found in covert and overt hearing suggested a similar neural code in the two states, which is also consistent with the findings of fMRI studies showing a similar neural substrate mediating auditory perception and imagery ^68–70^. It is also worth mentioning that the activation of the auditory cortex is not specific to speech imagery, as a recent study found simlilar response patterns also during music perception and imagery^71^. While these studues have established the feasibility of speech decoding in covert speech perception and production, further research is needed to devise system architectures and training procedures that can optimally fine-tune a model to perform and generalize well in both overt and covert conditions. Furthermore, expanding from the closed-set intelligible speech in this work to continuous, open-set, natural intelligible speech requires additional research, which will undoubtedly benefit from a larger amount of training data, higher-resolution neural recording technologies^72^, and the adaptation of regression models^73^ and the subject to improve the BCI system^26,27^.

In summary, we present a general framework that can be used for speech neuroprothesis technologies that can result in accurate and intelligible reconstructed speech from the human auditory cortex. Our approach takes a step toward the next generation of human-computer interaction systems and more natural communication channels for patients suffering from paralysis and locked-in syndromes.

**Figure 5.**
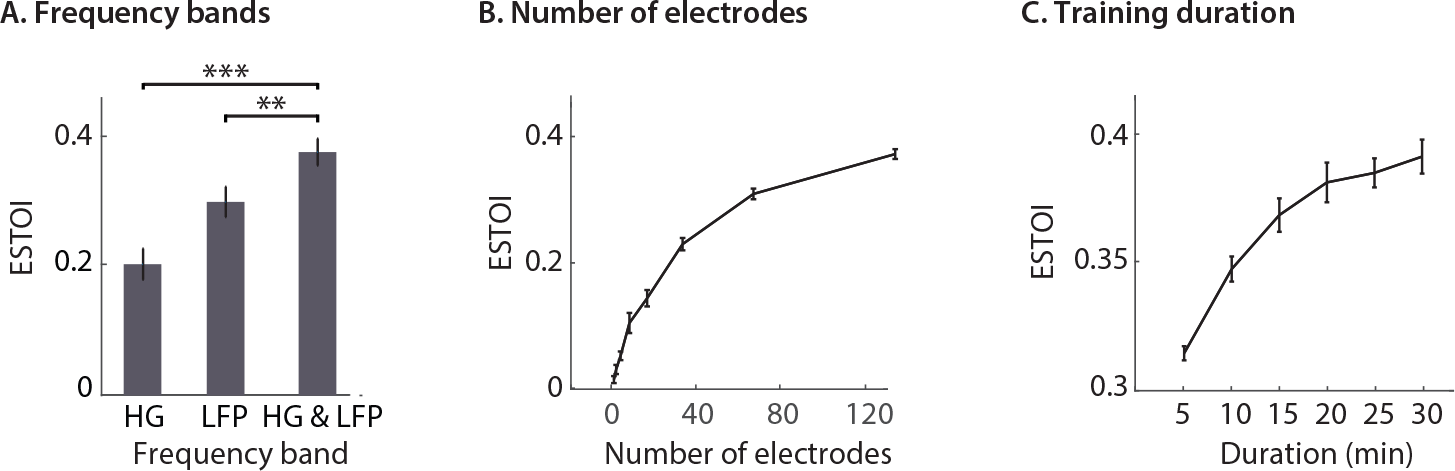
Effect of neural frequency range, number of electrodes, and stimulus duration on reconstruction accuracy. (A) The reconstruction ESTOI score based on high gamma, low frequency, and high gamma and low frequency combined. (B) The accuracy of reconstruction when the number of electrodes increases from one to 128. For each condition, 20 random subsets were chosen. (C) The accuracy of reconstruction when the duration of the training data increases. Each condition is the average of 20 random subsets.

## Materials and methods

### Participants and neural recording

Five patients with pharmacoresistent focal epilepsy were included in this study. All subjects underwent chronic intracranial encephalography (iEEG) monitoring at Northshore University Hospital to identify epileptogenic foci in the brain for later removal. Three subjects were implanted with only stereo-electroencephalographic (sEEG) depth arrays, one with a high-density grid, and one with both grid and depth electrodes (PMT, Chanhassen, MN, USA). The electrodes showing any sign of abnormal epileptiform discharges, as identified in the epileptologists’ clinical reports, were excluded from the analysis. All included iEEG time series were manually inspected for signal quality and were free from interictal spikes. All research protocols were approved and monitored by the institutional review board at the Feinstein Institute for Medical Research, and informed written consent to participate in the research studies was obtained from each subject before electrode implantation. All research was performed in accordance with relevant guidelines and regulations.

Intracranial EEG (iEEG) signals were acquired continuously at 3 kHz per channel (16-bit precision, range ± 8 mV, DC) using a data acquisition module (Tucker-Davis Technologies, Alachua, FL, USA). Either subdural or skull electrodes were used as references, as dictated by recording quality at the bedside after online visualization of the spectrogram of the signal. Speech signals were recorded simultaneously with the iEEG for subsequent offline analysis. Two ranges of neural frequencies were used in the study. Low-frequency (0-50 Hz) components of the neural data were extracted by filtering the neural signals using an FIR lowpass filter. The high-gamma (70-150 Hz) envelope^36^ was extracted by first filtering the data into eight frequency bands between 70 and 150 Hz using IIR filters. The envelope of each band was then obtained using a Hilbert transform. We took the average of envelopes in all frequency bands as the total envelope which was then resampled to 100 Hz. The high-gamma responses were normalized based on the responses recorded during a 2-minute silence interval before each recording.

### Brain maps

The electrode positions were mapped to brain anatomy using registration of the post-implant computed tomography (CT) to the pre-implant MRI via the post-op MRI^74^. After coregistration, the electrodes were identified on the post-implantation CT scan using BioImage Suite^75^. Following coregistration, the subdural grid and strip electrodes were snapped to the closest point on the reconstructed brain surface of the pre-implantation MRI. We used the FreeSurfer automated cortical parcellation^76^ to identify the anatomical regions in which each electrode contact was located within approximately 3 mm resolution (the maximum parcellation error of a given electrode to a parcellated area was < 5 voxels/mm). We used Destrieux’s parcellation because it provides higher specificity in the ventral and lateral aspects of the medial lobe^77^. The automated parcellation results for each electrode were closely inspected by a neurosurgeon using the patient’s coregistered post-implant MRI.

### Stimulus

The speech materials included continuous speech stories recorded in-house by four voice actors and actresses (duration: 30 min, 11,025 Hz sampling rate). Eight of the sentences (40 seconds) were used for objective tests and were presented to the patients eight times to improve the signal to noise ratio. The digit sounds were taken from the TI-46 corpus^35^. Two female (f2 and f8) and two male (m2 and m5) speakers were chosen from the corpus, and one token per digit and speaker was used (total of 40 unique tokens). Each digit was repeated six times to improve the signal to noise ratio of the neural responses. The speakers that uttered the digits were different from the speakers that narrated the stories.

### Acoustic representation

The auditory spectrogram representation of speech was calculated from a model of the peripheral auditory system^78^. The model consists of three stages: 1) a cochlear filter bank consisting of 128 constant-Q filters equally spaced on a logarithmic axis, 2) a hair cell stage consisting of a low-pass filter and a nonlinear compression function, and 3) a lateral inhibitory network, consisting of a first-order derivative along the spectral axis. Finally, the envelope of each frequency band was calculated to obtain a time-frequency representation simulating the pattern of activity on the auditory nerve^78^. The final auditory spectrogram has a sampling frequency of 100 Hz. The audio signal was reconstructed from the auditory spectrogram using an iterative convex optimization procedure^43^. For the vocoder-based speech synthesizer, we used the WORLD^45^ (D4C edition) system. In this model, four major speech parameters were estimated, from which the speech waveform was synthesized: 1) spectral envelope, 2) *f*0 or fundamental frequency, 3) band aperiodicity, and 4) voiced-unvoiced (VUV) excitation label. The dimension of each parameter was automatically calculated by the vocoder method and was based on the window size and the sampling frequency of the waveform (16 KHz).

### DNN architecture

We used a common deep neural network architecture that consists of two stages: feature extraction and feature summation^38–40^ (Fig. 2A). In this framework, a high-dimensional representation of the input is first calculated (feature extraction), which is then used to regress the output of the model (feature summation). The feature summation and feature extraction networks are optimized jointly together during the training phase. In all models examined, the feature summation step consisted of a two-layer fully connected network (FCN) with L2 regularization, dropout^41^, batch normalization^42^, and nonlinearity in each layer.

We study five different architectures for the feature extraction part of the network: the fully connected network (FCN, also known as the multilayer perceptron or MLP), the locally connected network (LCN) ^79^, convolutional neural network (CNN) ^80^, FCN+CNN, and FCN+LCN (for details of each architecture see Supp. Table 1). In the combined networks, we concatenated the output of two parallel paths, which were fed into the summation network. For FCN, the windowed neural responses were flattened and fed to a multilayer FCN. However, in LCN and CNN, all the extracted features were of the same size as the input, meaning that we did not use flattening, strided convolution, or downsampling prior to the input layer or between the two consecutive layers. Instead, the final output of the multilayer LCN or CNN was flattened prior to feeding the output into the feature summation network.

The optimal network structure was found separately for the auditory spectrogram and vocoder parameters using an ablation study. For auditory spectrogram reconstruction, we directly regressed the 128 frequency bands using a multilayer FCN model for feature extraction (Supp. Table 5). This architecture, however, was not plausible for reconstructing vocoder parameters due to the high-dimensionality and statistical variability of the vocoder parameters. To remedy this, we used a deep autoencoder network (AEC) ^47^ to find a compact representation of the 516-dimensional vocoder parameters (consisting of 513 spectral envelopes, pitch, voiced-unvoiced, and band periodicity) ^45^. We confirmed that decoding the AEC features performed significantly better than decoding the vocoder parameters directly (Supp. Table 2). The structure for the proposed deep AEC is illustrated in Figure 2D. To carry out decoding, we used a multilayer FCN, in which the number of the nodes changed in a descending (encoder) and then ascending order (decoder) (Fig. 2C)(supp. Table 6). The bottleneck layer of such a network (or the output of the encoder part of the pre-trained AEC) can be used as a low¬dimensional reconstruction target by employing the neural network model, from which the vocoder parameters can be estimated using the decoder part of the AEC. We chose the number of nodes in the bottleneck layer to be 256, because it maximized both the objective reconstruction accuracy (Supp. Table 3), and the subjective assessment of the reconstructed sound. To increase the robustness to unwanted variations in the encoded features, we used two methods in the bottleneck layer: 1) the hyperbolic tangent function (tanh) was used as a nonlinearity to control the range of the encoded features, and 2)

Gaussian noise was added during training prior to feeding into the first layer of the decoder part to make the decoder robust enough to unwanted changes in amplitude resulting from noises in neural responses. We confirmed that using additive Gaussian noise in the bottleneck instead of dropout performed significantly better (paired t-test, p<0.001). It is important that we use the same nonlinearity as the bottleneck (tanh) in the output of the main network, since the estimations should be in the same range and space as those in which they were originally coded. The best network architecture for decoding the vocoder parameters was found to be the FCN+LCN network (Supp. Table 4).

### DNN training and cross validation

The networks were implemented in Keras with a Tensorflow backend^81^. Initialization of the weights was performed using a previously proposed method which was specifically developed for deep multilayer networks with rectified linear units (ReLUs) as their nonlinearities^82^. It has been shown that using this method helps such networks converge faster. We used batch normalization ^42^, nonlinearity, and a dropout of p=0.3 ^41^ between each layer. We applied an *L2* penalty (with a multiplier weight set to 0.001) on the weights of all the layers in all types of networks (including the AEC). However, we found that using additive Gaussian noise in the bottleneck of the AEC instead of dropout and regularization performed significantly better (paired t-test, p<0.001). We used three types of nonlinearities in the networks: 1) LeakyReLU^83^ for all layers of AEC except the bottleneck and for all layers of the feature extraction part of the main network, 2) tanh for the output layer of the main network and the bottleneck of the AEC, and 3) the exponential linear unit (ELU) ^84^ for the feature summation network. Each epoch of training had a batch size of 256, and optimization was performed using Adam^85^ with an initial learning rate of 0.0001, which was reduced by a factor of two if the validation loss did not improve in four consecutive epochs. Network training was achieved in 150 epochs and was performed for each subject separately. The loss function was a combination of MSE and Pearson’s correlation coefficient for each sample:

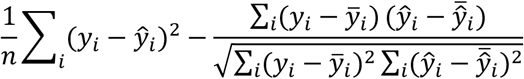

in which *y* is the actual label (auditory spectrogram or vocoder features) for that sample and 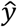 is the reconstruction from the output layer of the network. The maximum time-lag used was *τ*_max_ = 300 ms (Supp. Fig. 2). Because of the higher correlated activity between the neural responses of neighboring electrodes^86^, it was important to ensure that the networks can model the local structure in the data. Because both CNN and LCN use small receptive fields that take local patterns into account, we retained the spatial organization of the electrode sites in the input to the network, meaning that the electrodes that were close to each other in the brain were arranged to be close together in the input data matrix.

### Cross validation

We trained both the LR model and the DNN models using cross validation. We used the speech stories for training all models, and used repeated sentences (separate set from the stories) and digit sounds for testing. No digit sound was included in the training, and the speakers that uttered the digits were different from those that read the stories. The autoencoder network (AEC) was trained on a separate speech corpus (Wall Street Journal, WSJ, 80 hours of read speech) ^48^.

### Subjective and objective evaluations

We assessed the intelligibility of the reconstructed speech using both subjective and objective tests. For subjective assessment, 11 participants with self-reported normal hearing listened to the reconstructed digits using headphones in a quiet environment. Each participant listened to 160 tokens including 10 digits, four speakers, and four models. The participants were asked to report the digit or to select unsure if the digit was not intelligible. In addition, the participants reported the quality of the reconstructed speech using a mean opinion score (MOS): 1 (bad), 2 (poor), 3 (fair), 4 (good), and 5 (excellent). The participants also reported the gender of the speaker. For objective evaluation, we used the ESTOI measure^51^ which is a monaural intelligibility prediction algorithm commonly used in speech enhancement and synthesis research. The range of ESTOI measure is between zero (worst) and one (best).

## Data availability

The data that support the findings of this study are available upon request from the corresponding author [NM].

## Code availability

The codes for performing phoneme analysis, calculating high-gamma envelope, and reconstructing the auditory spectrogram are available at http://naplab.ee.Columbia.edu/naplib.html ^87^.

## Acknowledgments

We thank James O’Sullivan for providing helpful comments on the manuscript. This work was funded by a grant from the National Institutes of Health, NIDCD, DC014279 and the Pew Charitable Trusts, Pew Biomedical Scholars Program.

## Author Contributions

H. A., B.K., N.M., designed the experiment, evaluated the results and wrote the manuscript. B.K., J.H., N.M., A.M. collected the data. All authors commented on the manuscript.

